# Ether anesthetics prevents touch-induced trigger hair calcium-electrical signals excite the Venus flytrap

**DOI:** 10.1101/2021.08.18.456873

**Authors:** Sönke Scherzer, Shouguang Huang, Anda Iosip, Ines Fuchs, Ken Yokawa, Khaled A. S. AL-Rasheid, Manfred Heckmann, Rainer Hedrich

## Abstract

Plants do not have neurons but operate transmembrane ion channels and can get electrical excited by physical and chemical clues. Among them the Venus flytrap is characterized by its peculiar hapto-electric signaling. When insects collide with trigger hairs emerging the trap inner surface, the mechanical stimulus within the mechanosensory organ is translated into a calcium signal and an action potential (AP). Here we asked how the Ca^2+^ wave and AP is initiated in the trigger hair and how it is feed into systemic trap calcium-electrical networks. When *Dionaea muscipula* trigger hairs matures and develop hapto-electric excitability the mechanosensitive anion channel DmMSL10/FLYC1 and voltage dependent SKOR type Shaker K^+^ channel are expressed in the sheering stress sensitive podium. The podium of the trigger hair is interface to the flytrap’s prey capture and processing networks. In the excitable state touch stimulation of the trigger hair evokes a rise in the podium Ca^2+^ first and before the calcium signal together with an action potential travel all over the trap surface. In search for podium ion channels and pumps mediating touch induced Ca^2+^ transients, we, in mature trigger hairs firing fast Ca^2+^ signals and APs, found OSCA1.7 and GLR3.6 type Ca^2+^ channels and ACA2/10 Ca^2+^ pumps specifically expressed in the podium. Like trigger hair stimulation, glutamate application to the trap directly evoked a propagating Ca^2+^ and electrical event. Given that anesthetics affect K^+^ channels and glutamate receptors in the animal system we exposed flytraps to an ether atmosphere. As result propagation of touch and glutamate induced Ca^2+^ and AP long-distance signaling got suppressed, while the trap completely recovered excitability when ether was replaced by fresh air. In line with ether targeting a calcium channel addressing a Ca^2+^ activated anion channel the AP amplitude declined before the electrical signal ceased completely. Ether in the mechanosensory organ did neither prevent the touch induction of a calcium signal nor this post stimulus decay. This finding indicates that ether prevents the touch activated, glr3.6 expressing base of the trigger hair to excite the capture organ.

## Introduction

Within the plant kingdom the action potential (AP) of the carnivorous Venus flytrap is most similar to the all or nothing AP in our nerve cells (Hedrich and Neher, 2018). While APs in Arabidopsis and most other plants last >1 min and cannot be evoked repeatedly (Mousavi et al., 2013), the flytrap equivalent takes just about a second and can be fired a maximal frequency of 1 Hz (Bohm et al., 2016b). Compared to nerves with the information encoded by the frequency (number of APs per time), the flytrap system is more remote. It, however, allows *Dionaea* count to five APs (Bohm et al., 2016a). When a potential prey visiting the trap, attracted by color and odor, touches one of the six trigger hairs, an AP gets fired. The trigger hair is a mechano-sensitive organ that gets excited by share stress of >3° bending of 29 μN force (Scherzer et al., 2019). Two AP make the trap close and imprisons the animal prey. Three and more APs trigger touch hormone jasmonate (JA) signaling and JA production. A count of 5 is required to produce digestive enzymes and transporters that are specialist to take in the animal derived nutrients (Bohm et al., 2016a).

Hapto-electric energy conversion takes place in the indentation zone of the trigger hair podium. In this zone mechano-sensitive channels of the DmMSL10 and a DmOSCA1 type are expressed (Procko et al., 2021, Iosip et al., 2020)). MSL10 type channels in Arabidopsis and *Dionaea* operate as anion channels that upon activation depolarize cells (Maksaev and Haswell, 2012), while OSCAs rather Ca^2+^ (Yuan et al., 2014, Murthy et al., 2018, Thor et al., 2020). To trigger excitation in the flytrap cells have to be depolarized by 20-40 mV from the resting state (Hedrich and Fukushima, 2021). Thus, together MSL10 and OSCA(s) likely give rise to depolarization and Ca^2+^ influx for activation of anion channels. Anion channels in plants are the master switches in plant electrical excitation. The opening of anion channels and thus membrane electrical excitation is ultimately linked to the amplitude and kinetics of the given stimulus induced Ca^2+^ transient. Initial notice of a calcium-voltage-change association was made in the context of the wound/foraging response in the model plant Arabidopsis. Upon wounding the signaling molecule glutamate gets released to the extracellular space (Toyota et al., 2018), faced by ligand-binding site of GLRs (Gangwar et al., 2020, Alfieri et al., 2020). The Arabidopsis genome harbors 20 GLRs. Edgar Spalding lab in 2006 was first to demonstrate that glutamate evokes a Ca^2+^ signal as well as membrane depolarization which is suppressed when GLR3.3 is mutated (Qi et al., 2006). The Ted Farmer and Simon Gilroy labs have unequivocally shown that the electrical and Ca^2+^ signal, moving from the injured side in the local leaf to those wired to it via the vasculature, requires the presence GLR3.3 and 3.6 two glutamate receptors (Mousavi et al., 2013, Toyota et al., 2018).

Andrej Pavlovič and colleagues very recently have documented that ether suppresses AP firing and trap closing in *Dionaea* (Pavlovic et al., 2020). Cation channels responsible for the excitability of nerve cells are targets of anesthetics such as ether (Covarrubias et al., 2015, Brosnan and Pham, 2018). Key for anesthetics susceptibility are channel sites interacting with membrane lipids (Pavel et al., 2020).

Here we asked the flytrap ion channel target of ether. We found ether to suppress Ca^2+^ and AP stimulated trigger hairs excite capture organ. The fact that DmGLR3.6 expression was associated with flytrap excitability and the anesthetics suppressed glutamate-induced signals too, pointed to GLRs as likely ether targets.

## Results

### Calcium-electric signal originates in trigger hair podium

Our ecent studies have shown that the *Dionaea* trigger hairs can operate as sender and receiver of touch induced action potentials and Ca^2+^ waves (Iosip et al., 2020, Suda et al., 2020). How is this possible? The flytrap excitable cells rest around −140 to −120 mV. Whenever the membrane potential is depolarized below −100 mV an action potential is evoked.

For technical reasons one cannot measure APs in the very trigger hair and mechanical stimulate it at the same time. However, when voltage-recording microelectrodes were inserted in to the oblong mechano-sensitive cells of the podium of a receiver trigger hair, we could receive the AP induced by touch stimulation of one the other two mechanosensory organs operating as senders (Fig. S1).

In contrast to the AP the Ca^2+^ signal can be recorded in the same touch stimulated organ. Two types of Ca^2+^ signals were picked up: i) upon sub-threshold trigger hair bending the podium cells emitting GCaMP6f fluorescence first were those of indentation zone prone to sheering stress, while ii) above-threshold stimulations lighted up the entire podium that gave rise to selfsustained Ca^2+^ signal that left the trigger hair and propagated along the trap surface (Fig 1A and Supplementary Movie S1). In response to touch activation of the very trigger hair the podia of the other two touch sensitive organs of the same trap lobe lighted up as well (Supplementary Movie S2). Following the touch induced Ca^2+^ rise in the podium of the sender it dropped to base line levels with double exponential kinetics (tau_1_ = 0.55 ± 0.44 s and tau_2_ = 4.3 ± 2.02 s). The calcium fluorescence faded away in the indentation zone first, a finding in agreement with the notion that cells that were activated initially are subject of bleach first. This Ca^2+^-signal in the sender trigger hair and two receivers moved with a velocity of 2.66 ± 0.18 mm/s (mean ± SD; n=4) from the podium toward the hair tip.

**Fig. 1.**
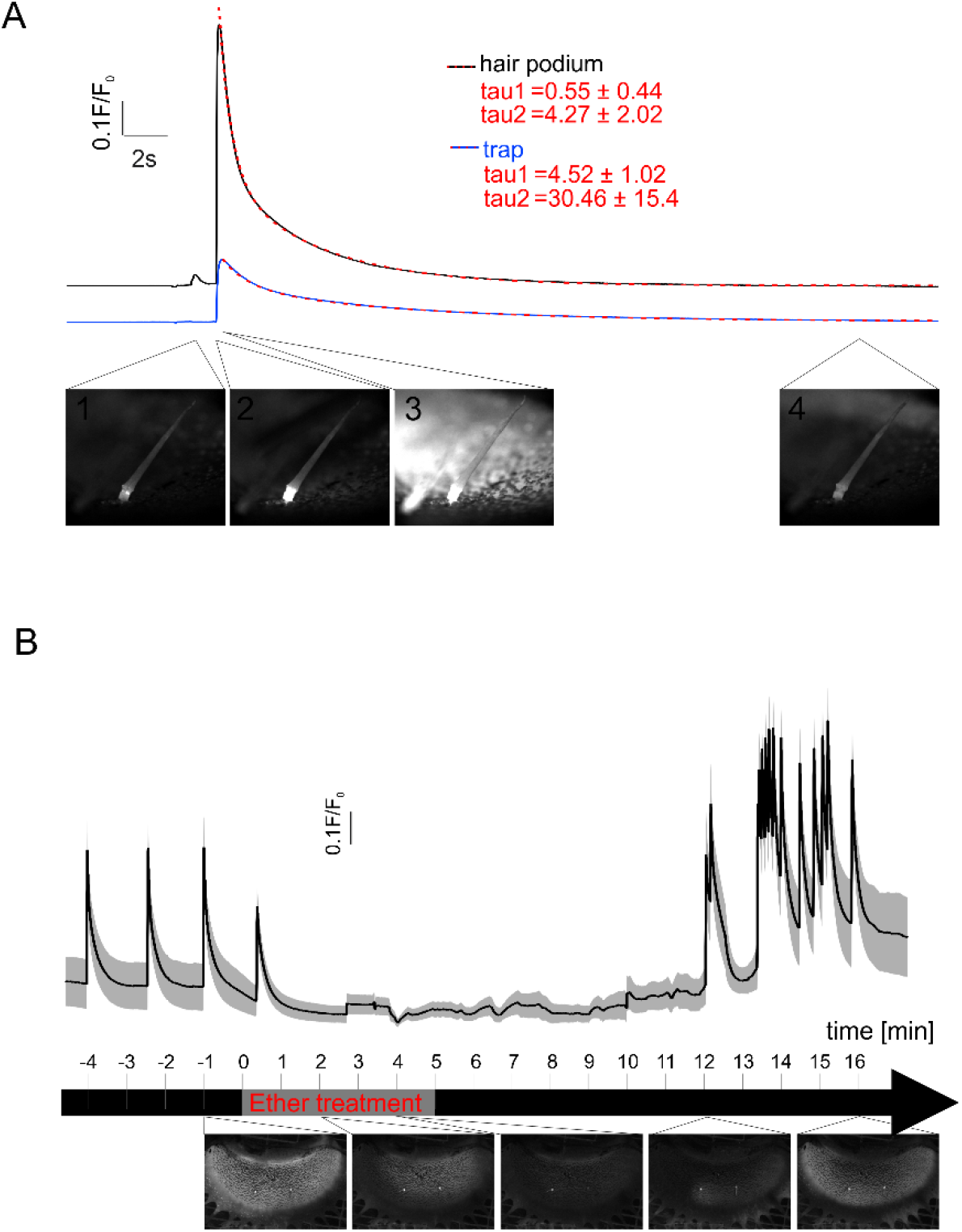
Ether inhibits hair podium [Ca]_ext_ spread into the trap. Representative time courses of the fluorescence intensity measured in the trigger hair podium (black) and in the trap (blue) as seen in supplementary movie S1. First subthreshold stimulus only was detected in the hair (image 1). The second suprathreshold stimulation occurred first in the trap podium (image 2) and then spread to the trap tissue (image3) before returning to baseline (image 4). Decay curves of fluorescence intensities were best fitted using a double exponential function (dotted red line) with the given tau values of 4 different traps (mean ± SD; n = 4). (B) Timeline of the ether anesthesia of a trap. The traps were mechanically stimulated on the trigger hair every minute. At timepoint 0, a saturated ether atmosphere was added for 5 minutes. After ventilation with fresh air, first Ca^2+^ signals were measured after 12 minutes again (mean ± SD (shading); n = 4). Below are sample images from the Supplementary Movie S3 of the indicated timepoints.

### Ether suppresses the touch induced Ca^2+^ wave

To investigate the ether effect on the Ca^2+^ signal, one trap lobe was cut off a *Dionaea* leaf and fixed a microscope stage. The sender trigger hair was stimulated every 1 min with ether fumigation initiated after the third AP (Fig1B and Supplementary Movie S3). Already after 2 min exposed to the ether atmosphere the touch-induced Ca^2+^-wave did not travel the entire trap anymore but just an area around the trigger hairs. In this situation the podia of the trigger hairs still emitted a pronounced GCaMP6f fluorescence. Another 2 min later, touch triggered a Ca^2+^-signal in the sender trigger hair only. When compared to ether-free conditions, we did not find ether to affect the nature and amplitude of the podium Ca^2+^ transient much. With prolonged anesthesia this situation did not change, indication that Ca^2+^ channels and Ca^2+^ ATPases (addressed below) involved in mechano-transduction and shaping the Ca^2+^ transient in the trigger hair podium do not present ether targets.

While replacing the ether atmosphere by fresh air it took about 7 min to see the touch induced Ca^2+^ wave coming back. Initially it did not travel the full trap surface, but the area right hand side of the capture organ displayed in the video (Fig1B and Supplementary Movie S3). Only 4 min later, however, the capacity of the trigger-hair induced Ca^2+^ wave had reconstituted fully.

### Ether suppresses the touch-induced AP

To study how ether affects the AP, we have placed a *Dionaea* plant in box that can be fumigated with ether. Traps were fixed to the recording chamber and a L-shaped glass capillary under the control of a micromanipulator positioned in proximity of a trigger hair. This computerized device allowed us to bend the trigger hair repeatedly and reproducibly (Supplementary Movie S4). Using surface electrodes initially, we recorded the AP in ether becoming progressively smaller until it faded away completely. In fresh air again the AP recovered its pre-stimulation shape (Fig. S2). To resolve how ether effect the AP different phases quantitatively, we inserted sharp intracellular electrodes into excitable lobe cells. In line with the literature, under control conditions the flytrap membrane potential depicted in Figure 2 was resting at about −120 mV (Böhm and Scherzer, 2021). Following trigger hair stimulation all-or-nothing APs got fired that under higher time resolution (insert in Fig. 2A) could be decomposed in to 6 phases (c.f. (Fabricant et al., 2021)): i) an initial fast depolarization phase to −60 mV which was followed by ii) a slower phase reaching the peak depolarization voltage of −20 mV. The depolarization phase is composed of iii) an initial fast component during which the membrane voltage repolarizes to −60 mV and iv) an initial lower component restoring the pre-stimulation voltage of −120 mV that merges with an even slower v) hyperpolarization phase that peaked at −130 mV. From this hyperpolarization overshoot the membrane potential in vi) final, a yet slow recovery phase returned to its pre-stimulus resting level again.

**Fig. 2.**
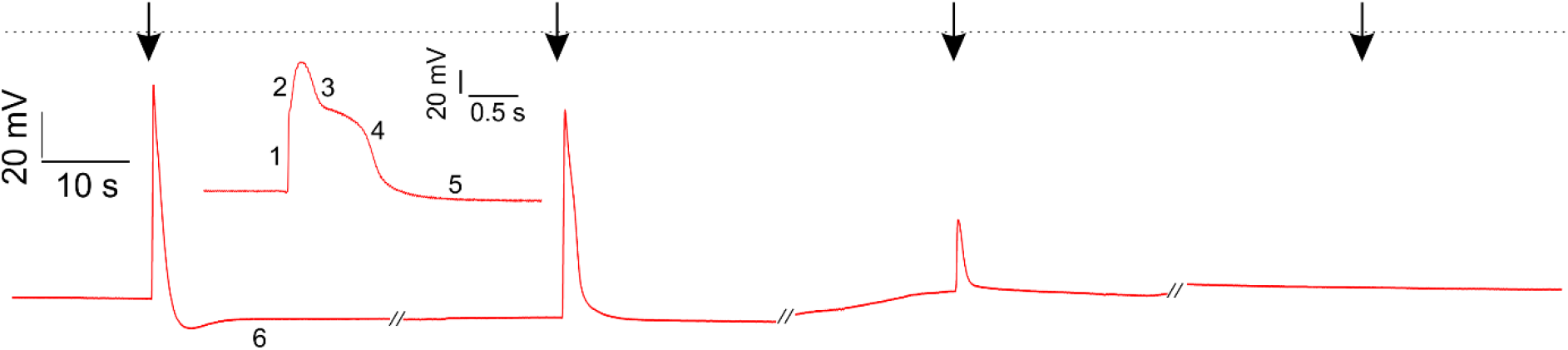
Ether inhibits electrical signal transduction in the traps. Dry impalement of a *Dionaea* trap. The magnification of the first AP illustrates the individual phases in the *Dionaea* AP. The trap was mechanically stimulated every 2 minutes (arrows) and a saturated ether atmosphere was added after the first AP. As a result, the resting potential remained relatively constant, while the depolarisation continued to decrease until no AP could be measured in the trap after 6 minutes.

When exposed to an ether atmosphere the resting potential did not change much but 5 min after anesthesia onset the peak depolarization dropped to – 40 mV and the overshoot hyperpolarization vanished completely. The overshoot results from a depolarization amplitude dependent transient hyperactivation of the AHA type H^+^ ATPase (Reyer et al., 2020). The depolarization phase of the *Dionaea* AP results from Ca^2+^ influx and Ca^2+^ activation of the anion channel (Hedrich and Fukushima, 2021, Beilby, 1984, Beilby, 2007, Shepherd et al., 2008). In presence of the general Ca^2+^ channel inhibitor La^3+^ touch induction both action potential (Hodick and Sievers, 1988) and Ca^2+^ wave is suppressed (Suda et al., 2020). This indicates that both signals are interconnected and that the Ca^2+^ channel blocker La^3+^ as well as ether anesthetics is suppressing the transition of Ca^2+^ signal and AP from the trigger hair podium into the trap. Anesthesia, however, does not seem to target podium located mechano-sensitive Ca^2+^ channels and Ca^2+^ ATPases (addressed below).

### Calcium channels and pumps located in the trigger hair podium

In Iosip et al. we have isolated trigger hairs and sequenced the RNAs expressed in the mechano-sensory organ (Iosip et al., 2020). Before trap had not opened during development it is regarded unmature and classified stage V (Fig. 3A and Fig. 4). We used stage V traps to compare touch/wounding calcium-electrical response. In contrast to mature traps in this developmental stage wounding did not evoke classical *Dionaea* APs but slow wave potentials (Fig. 3A). Upon touching trigger hairs with a fine brush, the entire unmature sensory organ and neighborhood co-stimulated accidentally lighted up (Fig. 3B and Supplementary Movie S5). The Ca^2+^ signal, however, remained local rather than traveling the stage V. This indicates that electrical and Ca^2+^ response exist in both stages but fast, long-distance calcium-electrical signal manifest only during maturation from stage V to VI.

**Fig. 3.**
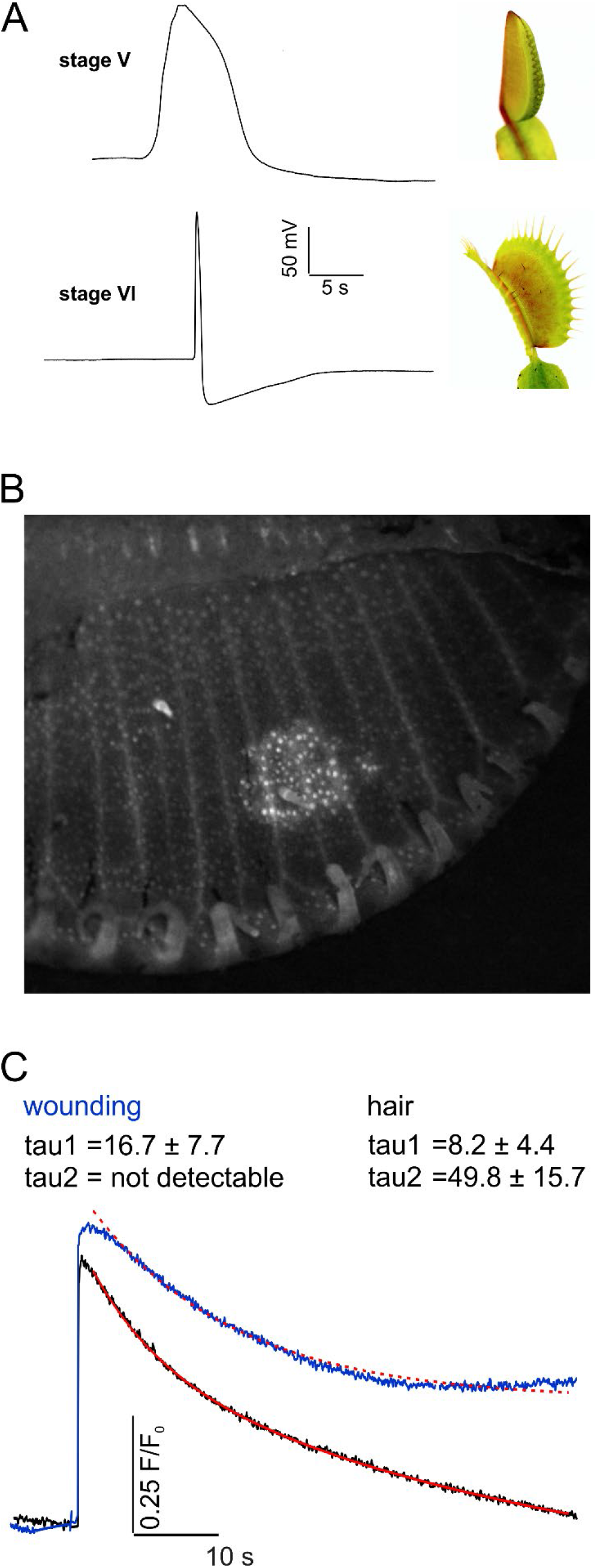
Wounding induced electrochemical signal transduction in *Dionaea* traps. Surface potential measurements of an immature stage V trap (top) and an adult stage VI trap (bottom) after wounding. Compared to the AP in adult traps, stage V traps only elicit a slow wave potential upon wounding. (B) The corresponding Ca^2+^ signal in stage V GCaMP6f traps is locally limited and does not spread fast over the whole trap (example image from Supplementary Movie S5). (C) Representative time courses of the fluorescence intensity measured in the wounded stage VI trap tissue (blue) and surrounding tissue (black) (from Supplementary Movie S6). The wounding results in a fast AP (c.f. (A)) which leads to a fast Ca^2+^ signal in the stage VI trap. However, unlike in the undamaged tissue, the signal does not decay rapidly in the wounded tissue. Thus, only the uninjured tissue can be well fitted with the double exponential function (red).

**Fig. 4.**
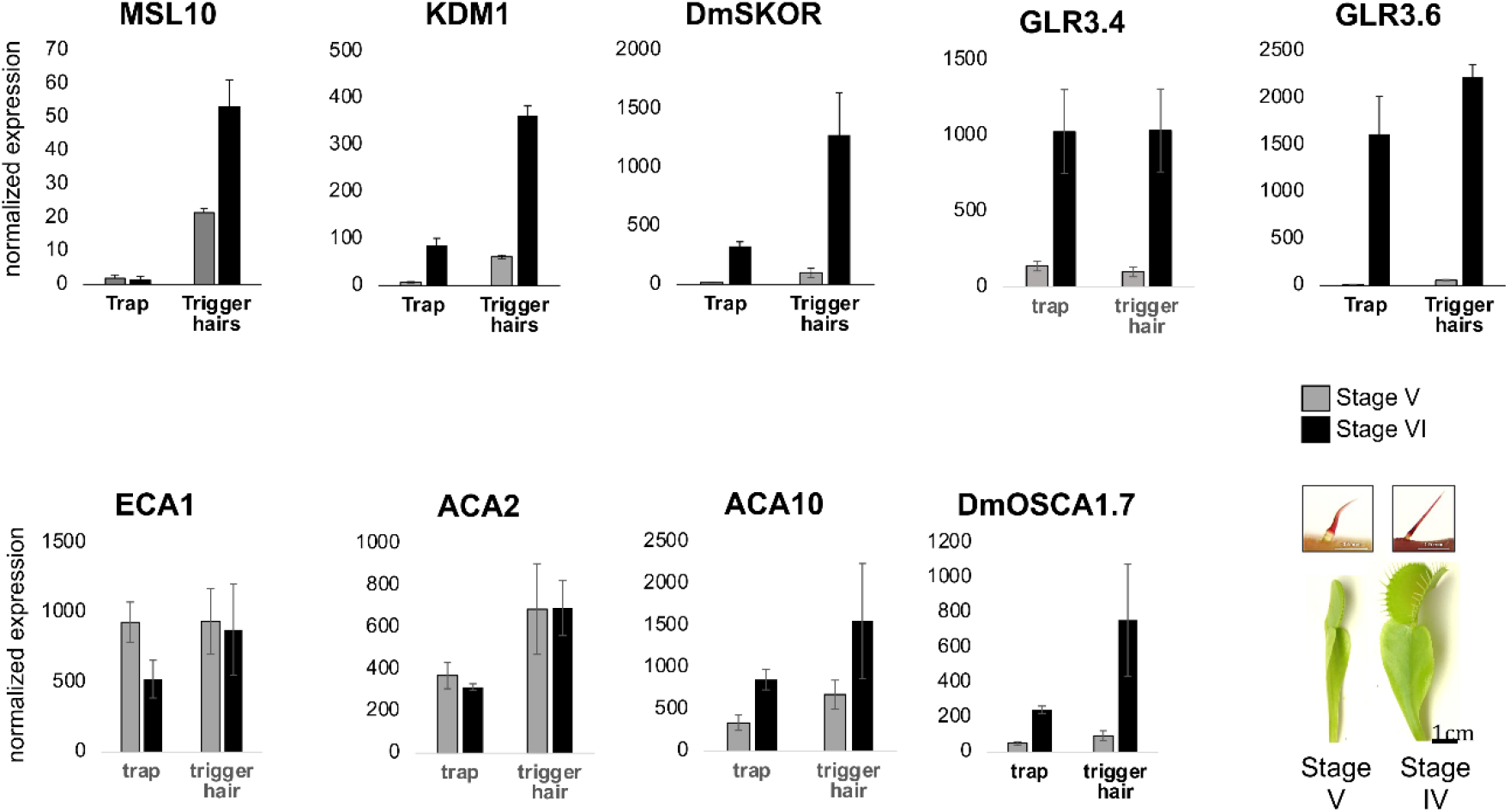
Developmental and tissue-specific expression analyses. Bottom right: Juvenile (stage V) trap with the corresponding juvenile TH and adult (stage VI) trap with the corresponding adult and electrically excitable TH. Normalized expression values of selected transporters (c.f. Supplementary Table S1) quantified by qPCR in stage V and stage VI trap and trigger hair (mean normalized to 10,000 actin ± SE; n = 3–6).

To spot the potential ether target, we reinvestigated and extended the RNA species that have been associated with Ca^2+^ channels, pumps, and carriers. The following search criteria were used: i) genes are expressed in the trigger hair (Iosip et al., 2020) ii) expression is induced when trigger hairs maturate, developing trigger hair-only mechano-sensitivity and iii) genes should predominately express in the podium.

Extensive transcriptomic analyses previously revealed the expression of numerous transporters in the *Dionaea* trigger hair (Iosip et al., 2020), with some being specific to the trigger hair such as MSL10 and the *Shaker* K^+^ channels KDM1 and DmSKOR. Moreover, the glutamate receptors DmGLR3.4 and DmGLR3.6 as well as the Ca^2+^-ATPase DmACA2 were characterized as trigger hair specific as well. Apart from DmGLR3.4 all of these transporters we found predominantly expressed in the podium fraction of mature trigger hair (Fig. 4, Supplementary Fig S4, and Table S1). The Ca^2+^-ATPase DmACA10 followed the same search criteria as well. qPCR analyses revealed that DmMSL10, KDM1, DmSKOR, DmGLR3.4, DmGLR3.6 and DmACA10 are transcriptionally induced in mature traps and fully functional stage VI trigger hairs, thus likely to contribute to the trap’s calcium and electrical excitability (Fig. 4). In a previous study, an OSCA (represented by Dm_00001755-RA in (Palfalvi et al., 2020)) was described as trigger hair specific. However, the expression of this gene is rather low. We therefore re-analyzed our trigger hair transcriptome and identified another OSCA1.7 type mechanosensitive Ca^2+^-channel in the trigger hair which we named Dm (Dm_00005287-RA) based on phylogenetic analyses. DmOSCA1.7 is expressed in the trigger hair and the mature trap and induced in both tissues during trap maturation and gain of electrical excitability (Fig. 4, Supplementary Fig S4, and Table S1).

In addition to DmOSCA1.7 we found DmGLR3.6 that represents homolog to Arabidopsis glutamate receptor channels GLR3.1/3.3 involved in long-distance mechanical stress electrical and Ca^2+^ signaling (Nguyen et al., 2018, Toyota et al., 2018). Loss-of-function mutants still respond to wounding locally but not systemically. In other words, in the mutant the slow wave potential (SWP; ~8 times longer and ~10 times slower compared to a *Dionaea* AP) and Ca^2+^ wave does not travel out of the locally wounded leaf. Furthermore, the Simon Gilroy lab has shown that upon wounding glutamate is released from injured cells (Toyota et al., 2018). When glutamate was applied to pre-wounded Arabidopsis leaves an electrical signal and a calcium wave was elicited (Toyota et al., 2018, Qi et al., 2006).

### Ether suppresses flytrap wounding and glutamate induced calcium signal transduction

Given glutamate release and GLRs are associated with wounding, we with an Eppendorf pipette tip pinched selected sites on traps (c.f. (Scherzer et al., 2019)) of GCaMP6f expressing *Dionaea* and monitored the calcium-electrical response. Following injury with a macroscopic pipette tip a rapid electrical- and calcium-signal was evoked that was spreading from the compressed trap site distally. In contrast to the stimulation of the trigger hairs (Supplementary Movie S6 left), the local calcium increase induced by the compression (right) was long-lasting. One min following pinch stimulation GCaMP6f fluorescence emission had drop by 49 % only (Fig. 3C and Supplementary Movie S6 right). When the trap was wounded by penetration with the pipette tip, the calcium signal in the vessels branching off from the injured section even outweighed the parenchyma signal (Supplementary Movie S7). The latter response might be explained by glutamate release in response to wound addressing GLR receptors to Ca^2+^ activate in the vasculature (c.f. (Toyota et al., 2018)). *Dionaea’s* response to glutamate was tested by applying 5 mM of the key amino acid in animal cells serving as neurotransmitter. To guarantee that extracellularly fed glutamate reach its targets in the trap, we cut off one trap lobe. After a wounding-recovery phase and the trap firing APs in response to trigger hair stimulation, glutamate was applied to the pre-wounded trap side (Supplementary Fig. S3). After adding the chemical stimulus to the vasculature-exposed midrib half, a fast AP and calcium wave travelled the trap (Fig 4B, Supplementary Movie S8).

To ask how ether is affecting the flytraps glutamate response, we initially stimulated traps via trigger hair bending once per minute (Supplementary Movie S9). After the third AP was fired, the trap was exposed an ether atmosphere. Five min later no trigger hair AP could be evoked. In this situation the trap was wounded. In this experimental series (Supplementary Movie S10) we analyzed GCaMP trap’s glutamate Ca^2+^ response in fresh air side-by-side with those exposed to ether. In the absence of ether, the touch-induced Ca^2+^ signal originating in trigger hair podium spread over the entire trap, while under anesthesia the touch induced Ca^2+^ rise was restricted to the very mechano-sensory organ (c.f. Supplementary Movie S3). When a 0.1 ml drop of 5 mM glutamate solution was applied in fresh air only it triggered a fast Ca^2+^ wave that moved from the side of midrib application all over the trap reaching epidermis cells, triggers hairs, nectaries, and very likely parenchyma cells too (Supplementary Movie S10).

This initial fast response was accompanied by a calcium wave travelling the vasculature. While the primary signals faded away the fluorescence of the latter became dominating. The more this kind of secondary Ca^2+^ wave propagated from the midrib towards the rim at a speed of only 0.3 ± 0.05 mm/s. It is worth mentioning that glutamate stimulation enlightened the vascular strands so that the entire venation pattern became visible: strands were running highly parallel in from midrib along the secretory gland zone, branched and fused with their neighbors in the nectaries zone, and grow extensions towards the teeth. In contrast ether treated traps did not respond to the chemical simulation with 5 mM glutamate. The application of 20 mM glutamate 2 minutes later had also no effect in the anesthetized trap. Together this indicates that with anesthetized traps touch Ca^2+^ signal is caught in the trigger hair podium. In this situation, long-distance Ca^2+^ signaling cannot be rescued by substituting mechanical stimulation by glutamate excitation.

## Discussion

### Trigger hair podium is side of hapto-calcium-electric signaling

We have shown that a threshold in the trigger hair displacement amplitude exists, that when passed gives rise to an AP (Scherzer et al., 2019) and a Ca^2+^ wave (Fig. 1A). Below this threshold the amount of ‘seed Ca^2+^’ entering the mechanosensitive cells of indentation zone does not auto-amplify the Ca^2+^ singnal. When the local Ca^2+^ rises above threshold, via a CICR (Calcium Induced Calcium Release)-related a self-propagating Ca^2+^ wave, it springs up in the indentation zone, ingresses the podium, and spreads all over the trap. We have documented that the podium differentially expresses DmOSCA1.7 a homolog of the mechanosensitive Ca^2+^ channel AtOSCA1 (Zhai et al., 2020, Yuan et al., 2014, Thor et al., 2020). In the working model (Fig. 5) Ca^2+^ influx via DmOSCA1.7 is triggering CICR via a yet unknown ER Ca^2+^ channel. The fact that Ca^2+^ signal propagates fast from podium to trap but slow from the podium to the trigger tip, points a apical-basal membrane asymmetry in Ca^2+^ transporters. Besides the OSCA the podium specifically expresses DmGLR3.6 type Ca^2+^ channel, which in Arabidopsis is required for a triggered local Ca^2+^ signal to become systemic (Nguyen et al., 2018, Toyota et al., 2018, Mousavi et al., 2013). In the model DmGLR3.6 is the Ca^2+^ window to the trap. The podium is giving birth to the Ca^2+^ as well as the AP. The AP is initiated by the opening of a depolarization-and Ca^2+^ activated anion channel such as DmQUAC1 (c.f. (Imes et al., 2013, Mumm et al., 2013, Dreyer et al., 2012)). Upon mechanical activation of DmMSL10 anions are released and pre-depolarize the membrane. This voltage and OSCA/CICR Ca^2+^ above threshold activation initiated the depolarizing phase of the AP. The Shaker type voltage (depolarization) dependent GORK/SKOR type K^+^ channel is preferentially expressed in the podium and engaged with the repolarization of the AP. For the membrane potential to recover the resting state, the QUAC1 type anion channel the cytoplasmic Ca^2+^ level must drop. This task is accomplished by the two podium differentially expressed plasma membrane Ca^2+^ ATPase DmACA2 and 10 the ER Ca^2+^ pump DmECA1.

**Fig. 5.**
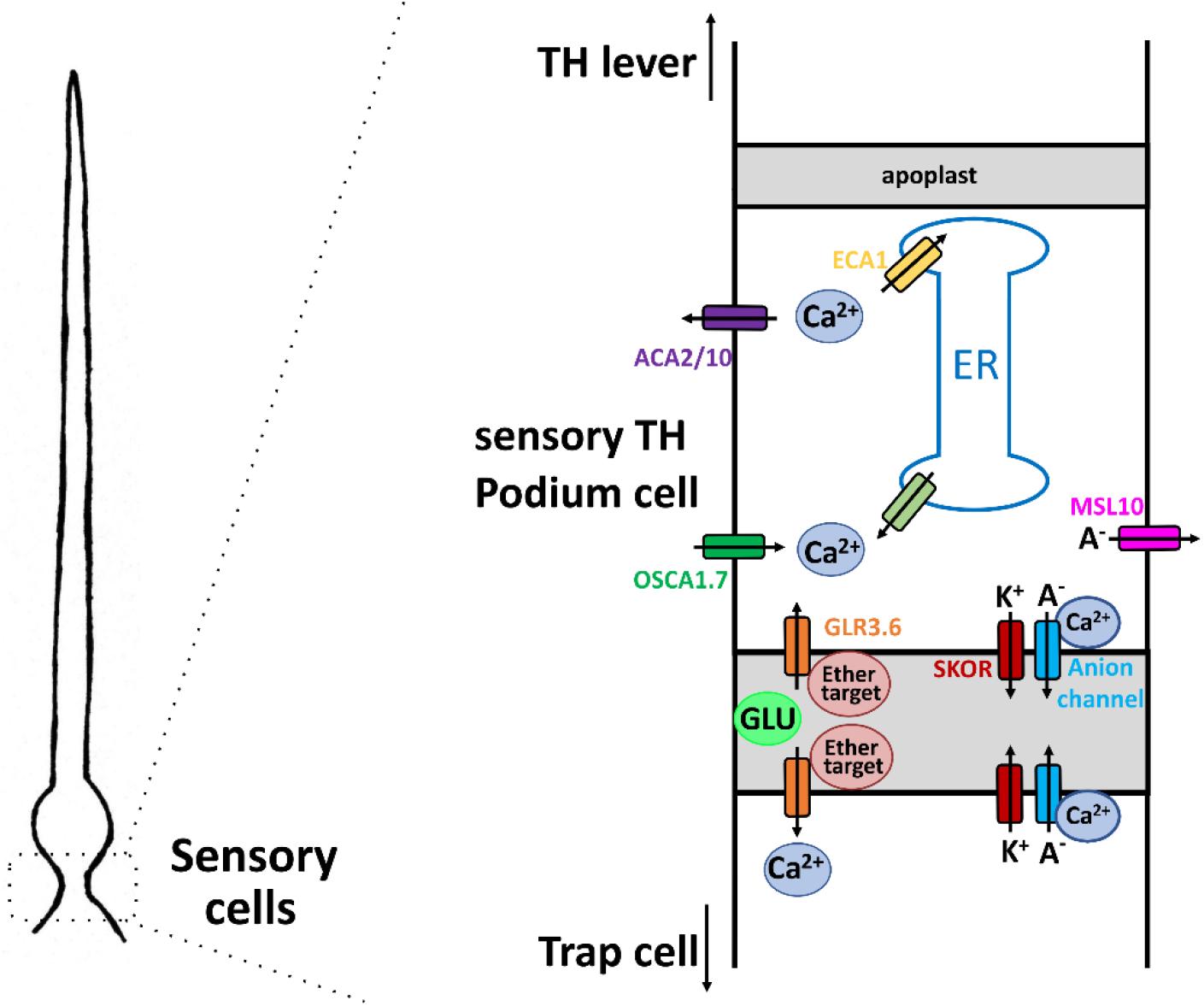
Working hypothesis on signal formation and propagation in the trigger hair. Schematic drawing of a trigger hair (left) and the magnification of a sensory podium cell (right). Mechanical stimulation activates MSL10 (pink) and OSCA1.7 (green) which results in a depolarization an increased [Ca^2+^]cyt, which further increases via calcium induced calcium release (light green) of the endoplasmic reticulum. Ca^2+^-induced activation of anion channels (blue) completes the depolarisation phase of the AP, which is repolarised by activation of potassium channels (red). The resting calcium concentration is restored by the ECA1 (yellow) and ACA2/10 (purple) pumps. Apoplastic glutamate activates GLR3.6 channels (orange) and transits the Ca^2+^ signal into the unstimulated trap. The application of ether (red) most likely suppresses Ca^2+^ signalling into the trap by blocking these glutamate receptors and thus the electrical AP in the trap remains absent.

### Ether counteracts glutamate induced calcium-electrics

The primary site of action of general anesthetics is the nervous system, where anesthetics like ether inhibit neuronal transmission. Anesthetics directly target a subset of plasma membrane cation channels. Our studies have shown that ether does not impair touch induced Ca^2+^ transients, excluding an OSCA as target of inhibition. Ether anesthetics and the Ca^2+^ channel blocker La^3+^ (Hodick and Sievers, 1988), however, prevents the touch- and glutamate induced Ca^2+^ electric signal to exit the podium. A similar lack of mechanical and glutamate dependent initiation of long-distance calcium electrical signal is known from an Arabidopsis mutant that lost glr3.1/3.3. These two AtGLRs were found to express in the vasculature (Toyota et al., 2018).

The *Dionaea* genome encodes 22 GLRs and DmGLR3.6 is differentially expressed in the trigger hair podium. Upon glutamate application the vasculature emitted a particularly strong GCAMP6f Ca^2+^ fluorescence emission but not in the presence of ether. It is thus tempting to speculate that in *Dionaea* DmGLR3.6 in the trigger hair podium and those expressed outside the mechano-sensor represent ether targets. This working hypothesis in future studies can be tested for DmGLR3.6 functionally complementation and anesthesia of the Arabidopsis glr3.1/3.3 mutant.

## Materials and methods

### Microscopy

Stereoscopic images and videos were taken using a Leica M165 FC fluorescence stereo microscope, a Leica EL6000 lightsource, the Leica filter set ET GFP LP (Leica Microsystems, Germany), and a cooled CCD camera (Hamamatsu C4742-80-12AG, Hamamatsu Photonics, Herrsching, Germany). For GCaMP6f calcium imaging, an excitation wavelength of 475 nm and an exposure time of 50 ms at 2x binning was used. Fluorescence videos were taken as a sequence of uncompressed tiff files in a range where the fluorescence intensities of samples and brightness values of captured images were linear. Ether was applied by adding 2 ml ether to a glass container next to the sample in a gas-tight setup.10x time laps for Movies S3, S8, S9 and S10 was achieved using Windows Movie Maker (Microsoft), Side by side views were produced in Premiere Pro (Adobe) and compression to MPEG2 was done in VLC media player (VideoLan). To calculate the propagation velocity of the [Ca^2+^]_cyt_ increase a sample rate of 25 ms was chosen. Hairs and traps were background subtracted and the mean velocity was calculated over known distances. For quantitative analysis of areas with increased [Ca^2+^]_cyt_ after stimulation two ROIs (regions of interest) with the same pixel numbers were defined in the indicated tissues using ImageJ. Data points were fitted using Igor Pro 8.

### Plant material and tissue sampling

*Dionaea muscipula* GCaMP6f plants were provided from Mitsuyasu Hasebe lab (Suda et al., 2020) and grown in plastic pots at 22 °C in a 16:8 h light:dark photoperiod. For microscopic experiments traps were bisected and the half which was still connected to the plant was fixed to a microscope slide the day before measurements.

For the qPCR expression measurements of genes of interest in wild-type plants, the trigger hair was dissected in two parts: the basal part and the tip. Entire/un-dissected trigger hairs were also collected as a control. For the whole trigger hairs qPCR expression analysis 300 - 600 trigger hairs were needed for one replicate, while for the tip and the base parts between 880 and 940 trigger hairs were needed to extract enough RNA for one replicate. Three replicates were used in total n = 3.

For qPCR analyses, RNA was isolated from each sample using the NucleoSpin Plant RNA extraction kit (Macherey-Nagel, Düren, Germany) according to the manufacturer’s instructions and in combination with Fruit-mate for RNA Purification solution (Takara Bio Europe SAS, Saint-Germain-en-Laye, France). Briefly, 100 mg of powdered plant material was thoroughly mixed with 350 μl of Fruit-mate solution for 1 minute. Following 10 minutes of centrifugation (20,000 rcf) at 4°C, the supernatant was mixed with the lysis buffer provided by the kit (RAP), together with 3.5 μl TCEP (Tris(2-carboxyethyl)phosphine hydrochloride, 0.5M, pH 7, Sigma-Aldrich). Apart from this, the kit manufacturer’s instructions were followed except for the DNA digestion, which was performed in a separate step after the RNA isolation. For 1 μg of RNA, 3 μl DNase Buffer (10x Reaction Buffer with MgCl2 for DNase, Thermo Fisher Scientific, Darmstadt, Germany), 0.5 μl DNase inhibitor (RiboLock RNase Inhibitor 40 U/μl, Thermo Fisher Scientific), and 1 μl DNase I (DNase I, RNase free 1000 units (1U/μl), Thermo Fisher Scientific) were mixed in a final volume of 30 μl and incubated at 37°C for 30 minutes. Next, the DNA-free RNA was precipitated in isopropanol overnight at −20°C, together with 1% glycogen (RNA Grade (20 mg/ml), Thermo Fisher Scientific), 10% NH4-Ac (5 mM in EDTA), 60% Isopropanol (2-Propanol AppliChem, Darmstadt, Germany), and water up to a final volume of 100 μl. The samples were washed using 70% Et-OH, centrifuged at 4°C for 20 minutes, and the resulting pellet was dried at 37°C and resuspended in water (DEPC, AppliChem).

The RNA was transcribed into cDNA using the M-MLV Reverse Transcriptase (RNase H- Point Mutant, Promega, Walldorf, Germany). qPCR was performed using a Realplex Mastercycler system (Eppendorf, Hamburg, Germany) and ABsolute QPCR SYBR green capillary mix (Thermo Fisher Scientific). Quantification of the actin transcript DmACT1 (GenBank: KC285589, Dm_00017292-RA) and transcripts for D. muscipula genes of interest was performed by real-time PCR. D. muscipula transcripts were normalized to 10,000 molecules of DmACT1.

### Membrane potential measurements

Before the dry-impalement experiments, a trap was cut in half and the half that was still connected to the plant was fixed in a Petri dish on a chlorinated silver wire as a reference electrode with double-sided adhesive tape. After a 30-minute recovery, the trap was inserted with a sharp impalement electrode. For impalements, microelectrodes from borosilicate glass capillaries with filament (Hilgenberg,) were pulled on a horizontal laser puller (P2000, Sutter Instruments) and filled with 300 mM KCl and connected via an Ag/AgCl half-cell to a headstage (1 GΩ, HS-2A, Axon Instruments). An IPA-2 amplifier (Applicable Electronics) was used, and the cells were impaled by an electronic micromanipulator (NC-30, Kleindiek Nanotechnik).

To anaesthetise the trap in Figure 1, 1 ml of ether was placed in a glass vial next to the trap and the trap was continuously stimulated at the trigger hair at 2-minute intervals until no more AP could be recorded.

To test the glutamate effect on the trap, 20 μl of 5 mM sodium L-glutamate was pipetted either onto the trap surface or onto the cut edge.

For surface potential measurements in Fig S2 and Supplementary Movie S4, we measured the extracellular potential of the trap tissue. One silver electrode was impaled into the trap surface. The reference electrode was put into the wet soil or the petiole. Electrical signals were amplified 100× and recorded with PatchMaster software (HEKA). Trigger hair stimulation or wounding to the trap tissue was applied at the given timepoints by a motorized arm.

## Acknowledgements

We thank Tracey Ann Cuin for her helpful discussions and assistance in preparing this paper. We would like to thank Christian Wegener (Professor of Neurogenetics Biocenter Würzburg) for his kind support and for making his setups available.

## Supporting information

### Supplemental Figures

**Fig. S1.**
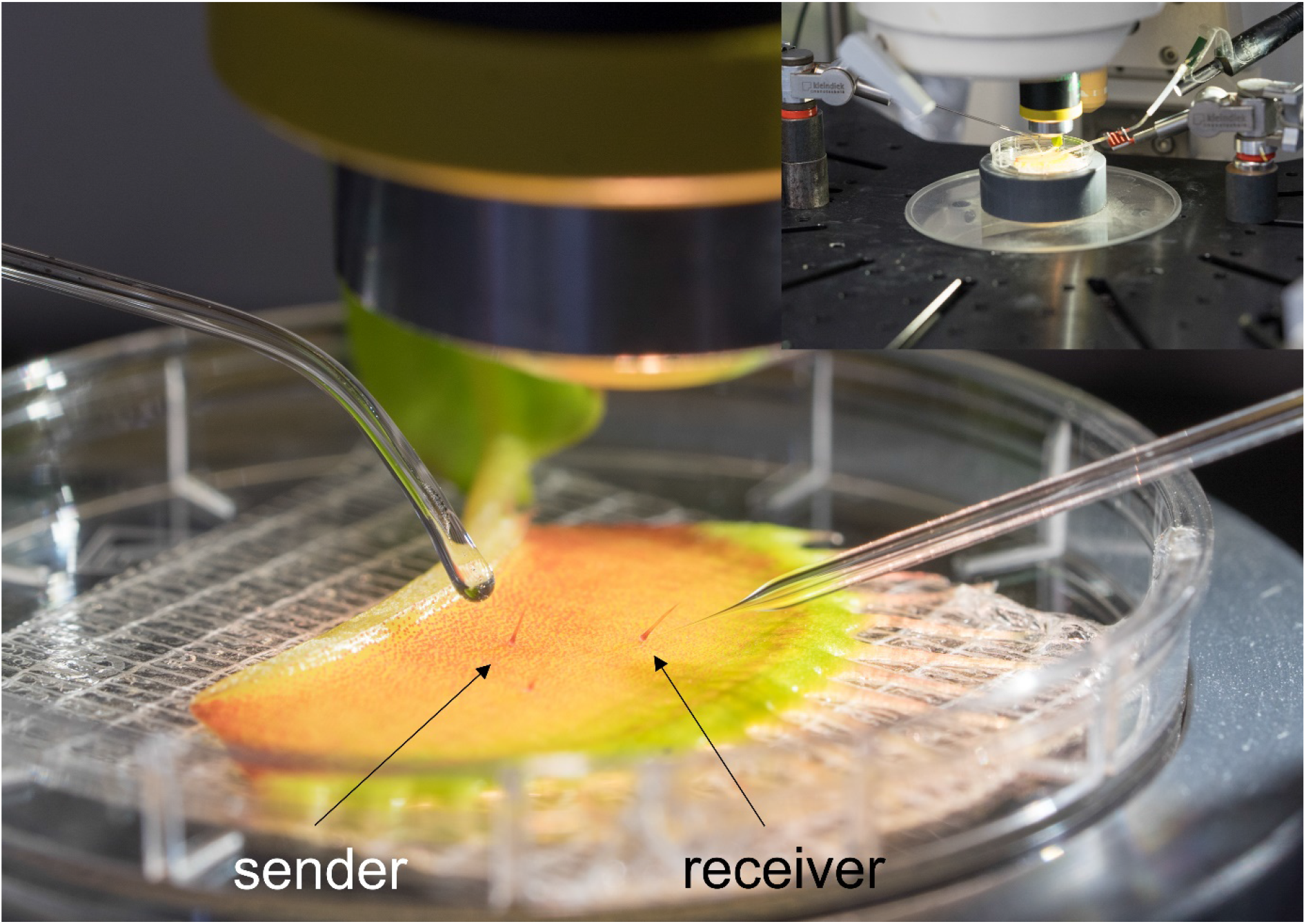
Exemplary setup of the voltage recording. The trap half is fixed under the microscope (upper right). While one trigger hair (sender) can be stimulated from the left, another hair (receiver) is impaled from the right.

**Fig. S2.**
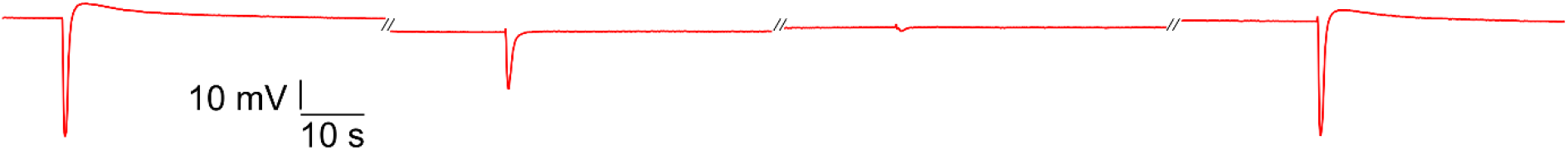
Representative surface potential measurement of a *Dionaea* trap under an ether atmosphere as seen in Supplementary Movie S4, with following ventilation.

**Fig. S3.**
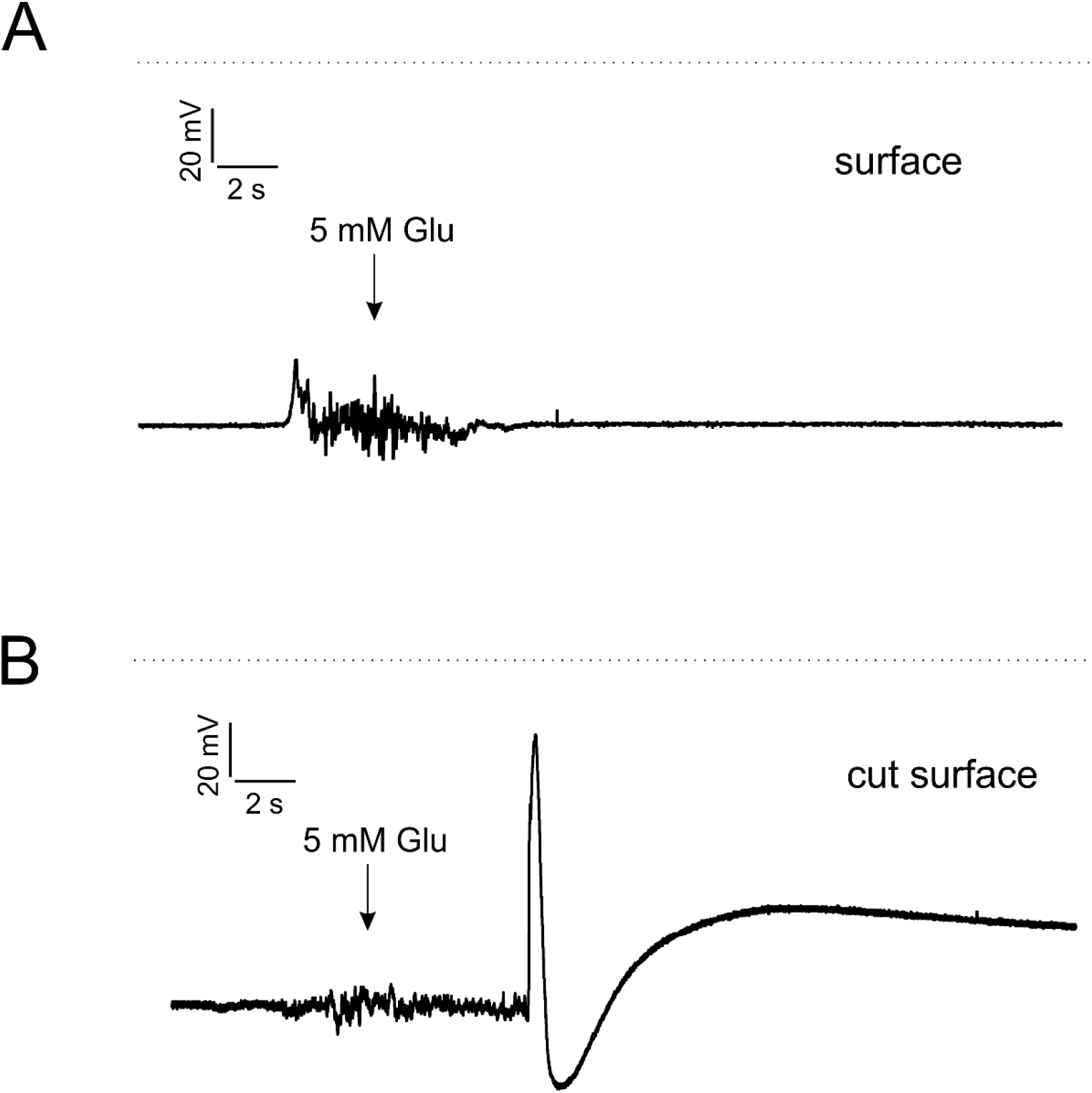
Membrane potential of a *Dionaea* trap where 5 mM glutamate (arrow) was applied to the undamaged surface (up) ore the cut surface (down). Only application to the cut surface initiated an AP.

**Fig. S4.**
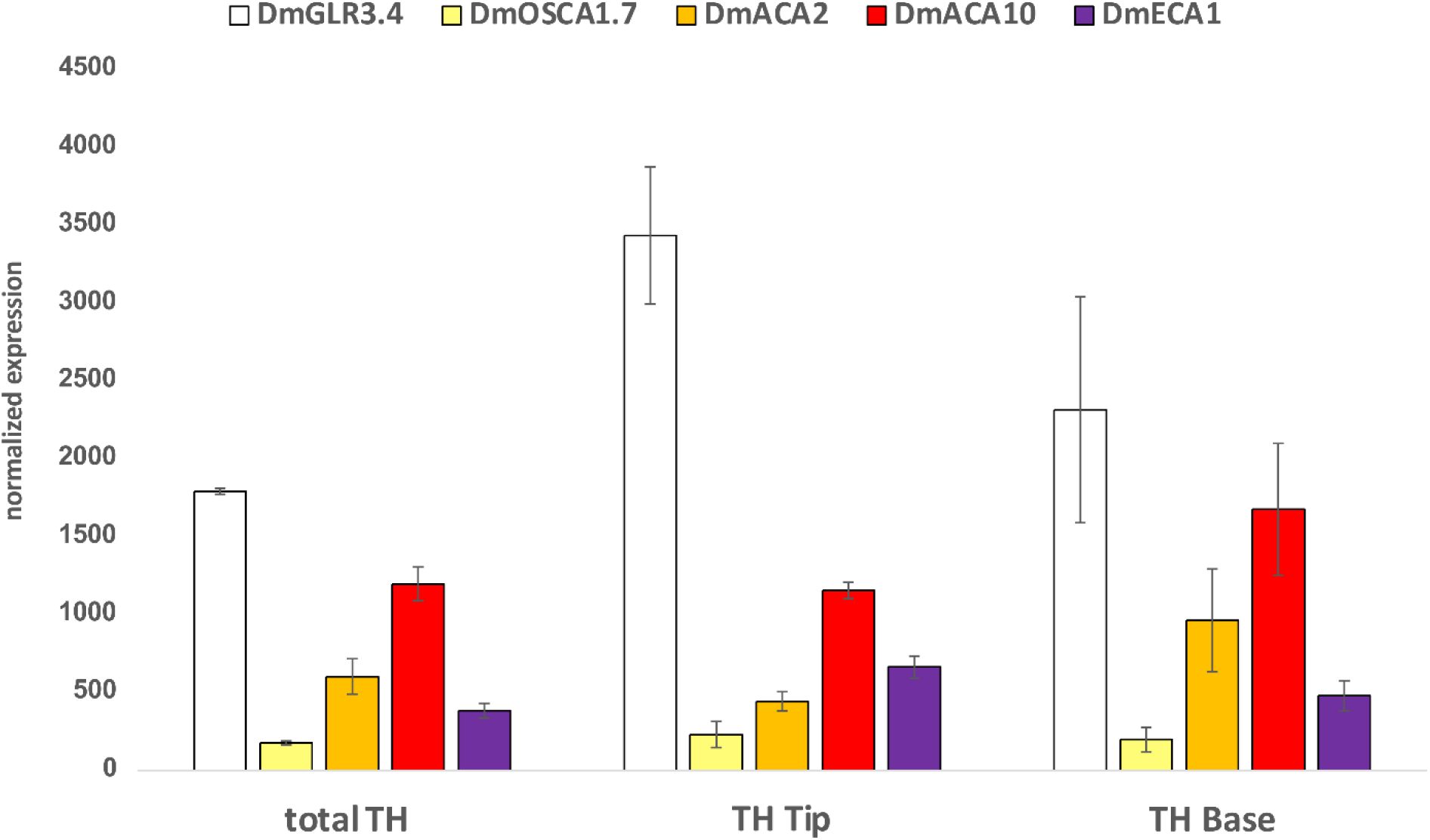
Normalized expression values of selected transporters quantified by qPCR in the whole trigger hair, in the tip/lever of the trigger hair, and in the base/podium of the trigger hair (mean normalized to 10,000 actin ± SE; n = 3).

**Supplemental Table S1.**

| Gene (ID) | Identifier | TH specific expression | Podium (enriched) expression | Enriched in trap stage V vs VI | Enriched in trigger hair stage V vs VI |
| --- | --- | --- | --- | --- | --- |
| **DmMSL10** | Dm_00009130-RA | + | + | - | (+) |
| **KDM1** | Dm_00004067-RA | + | + | + | + |
| **DmSKOR** | Dm_00007946-RA | + | + | + | + |
| **GLR3.4** | Dm_00004609-RA | + | - | + | + |
| **GLR3.6** | Dm_00002270-RA | + | + | + | + |
| **ECA1** | Dm_00014557-RA | - | - | - | - |
| **ACA2** | Dm_00018589-RA | + | + | - | - |
| **ACA10** | Dm_00009018-RA | - | + | (+) | (+) |
| **DmOSCA1.7** | Dm_00005287-RA | - | - | + | + |

### Supplemental Movie legends

Similar results were obtained from at least three biologically independent GCaMP6f expressing plants for all shown Supplementary movies.

**Movie S1**

Representative movie of a GCaMP6f expressing *Dionaea* trap where the trigger hair is mechanical stimulated. The first stimulus (6 s) was subthreshold, while the following (7 s) was initiating a Ca^2+^ signaling.

**Movie S2**

Young *Dionaea* traps without the characteristic pigmentation also show a Ca^2+^ signal in the digestive glands after mechanical stimulation.

**Movie S3**

10x time-lapse video of a *Dionaea* trap stimulated once per minute. After minute 3, a saturated ether atmosphere was created for 5 minutes, which was then replaced with fresh air. Please note that even under full anesthesia only the stimulated trigger hair podium shows an unaffected Ca^2+^ signal.

**Movie S4**

Setup used for surface potential measurements as presented in Supplementary Fig. S2.

**Movie S5**

Juvenile stage 5 trap mechanically stimulated with a soft brush. Please note that the Ca^2+^ signal does not spread over the whole trap but remains localized.

**Movie S6**

A *Dionaea* trap mechanically stimulated with a fine brush on the trigger hair on the left and pressure applied with a pipette tip on the right. Please note that both stimulations elicit a fast Ca^2+^ signaling in the entire trap, but the calcium signal at the site of the pressure decays very slowly.

**Movie S7**

Higher resolution of intense wounding leading to a particularly strong calcium signal in the vasculature.

**Movie S8**

10x time-lapse video of a trap where 5 mM glutamate was applied on the trap surface (3 s) not leading to any calcium signaling. Only application on the cut midrib (9 s) resulted in rapid and long-lasting calcium signaling, especially in the vasculature. As a control, a trigger hair was mechanically stimulated at 18 s, which resulted in a regular calcium signal.

**Movie S9**

10x time lapse of a trap that was mechanically stimulated once per minute. After 3 min, ether was added. When calcium signal transduction failed in the trap, strong injury was applied at 30 sec. Please note that again under anesthesia mechanical stimulation still resulted in a calcium signal in the podium and also the injury did not result in a calcium signal propagation in the trap.

**Movie S10**

10x time lapse of two *Dionaea* traps side by side. In both traps, a mechanical control AP was triggered at 6 s. Afterwards, only the right trap was treated with ether and after 8 minutes (54 s) a hair was stimulated again to verify that the right trap was anaesthetized. Subsequently (100 s), 5 mM glutamate was applied to the incised midrib in both traps, resulting in a Ca^2+^ signal only in the untreated left trap. Even a new application of 20 mM glutamate (72 s) to the anaesthetized trap did not lead to a calcium signal.

